# Mesoscale Explorer - Visual Exploration of Large-Scale Molecular Models

**DOI:** 10.1101/2024.09.02.610826

**Authors:** Alexander Rose, David Sehnal, David S. Goodsell, Ludovic Autin

**Affiliations:** independent; National Centre for Biomolecular Research, Faculty of Science, Masaryk University, 625 00, Brno, Czech Republic; Department of Integrative Structural and Computational Biology, The Scripps Research Institute, La Jolla, CA 92037, USA; Research Collaboratory for Structural Bioinformatics Protein Data Bank, Rutgers, The State University of New Jersey, Piscataway, NJ 08854, USA

**Keywords:** molecular graphism, mesoscale models, web-based 3D visualization, interactive tours, 3D animation

## Abstract

The advent of cryo-electron microscopy (cryo-EM) and cryo-electron tomography (cryo-ET), coupled with computational modeling, has enabled the creation of integrative 3D models of viruses, bacteria, and cellular organelles. These models, composed of thousands of macromolecules and billions of atoms, have historically posed significant challenges for manipulation and visualization without specialized molecular graphics tools and hardware. With the recent advancements in GPU rendering power and web browser capabilities, it is now feasible to render interactively large molecular scenes directly on the web. In this work, we introduce *Mesoscale Explorer*, a web application built using the *Mol\** framework, dedicated to the visualization of large-scale molecular models ranging from viruses to cell organelles. *Mesoscale Explorer* provides unprecedented access and insight into the molecular fabric of life, enhancing perception, streamlining exploration, and simplifying visualization of diverse data types, showcasing the intricate details of these models with unparalleled clarity.

Statement: *Mesoscale Explorer* leverages advanced GPU rendering and web technologies to facilitate and democratize the interactive 3D visualization of large-scale molecular models from viruses to cellular organelles composed of millions of atoms. *Mesoscale Explorer* enables broader exploration and deeper understanding of the complex structure of these large molecular landscapes.

## Introduction

In 1966, the first 16mm movies of animated proteins were produced by C. Levinthal^1^. This pioneering work was followed by a movie of the Insulin Dimer made by the *Molecular Modelling System*^2^ (MMS) devised by J. J. Beitch, R. A. Ellis, J. M. Fritch and G. R. Marshall. Since then, technological advancements have continually enhanced tools for interactively manipulating proteins and generating explanatory animations and movies^3,4^.

Several interactive molecular viewers have been developed over the years, including *GRAMPs*^5^, *Kineimage*^6^, *rasmol*^7^, *VMD*^8^, *Ribbon*^9^, *PMV*^10^, *Pymol*^11^, *BallView*^12^, *Chimera*^13,14^, and *YASARA*^15^. The advent of the internet has further democratized molecular structure manipulation with web-based viewers like *Jmol*^16^, *NGL*^17^, *LiteMol*^18^ and *iCn3D*^19^. As interactivity became more accessible, additional features for animation and movie making were developed in desktop applications (e.g., *YASARA* movies (https://yasara.org/movies.htm), *eMovie*^20^, *ChimeraX* movie^21^) as well as plugins for general 3d animation software (*mMaya* (https://clarafi.com/tools/mmaya/), *ePMV*^22^, *Molecular Nodes* (https://bradyajohnston.github.io/MolecularNodes/)) and web platforms (*PolyviewMM*^23^, *movieMaker*^24^, *PMG*^25^, *activeICM*^26^). This evolution led to the creation of outreach platforms such as *Proteopedia* (https://proteopedia.org), *FirstGlance* in *Jmol* (https://www.bioinformatics.org/firstglance/fgij/), and *Jolecule* (https://jolecule.appspot.com/), which provide guided macromolecular visualization with ease of use.

However, these platforms often struggle with the increasing complexity and size of contemporary molecular models, particularly with the advent of the “resolution revolution” in cryo-electron microscopy^27^ (cryo-EM and cryo-ET) and multiscale molecular modeling techniques such as *cellPACK*^28,29^, *YASARA* PETWORLD^30^, and *Mesocraft*^31^. These advances have enabled the creation of comprehensive 3D models of viruses, bacteria, and cellular organelles, comprising thousands of macromolecules and billions of atoms. The widespread availability of these models has opened new development avenues for their exploration^32–34^ and rendering^35–38^. New ways to visualize molecular structures in their cellular context are emerging, ranging from guided tours in Apple Vision Pro with *cellWalk* (https://cellwalk.ca/) to entire educational courses built around mesoscale animations in *SmartBiology* (https://www.smart-biology.com). It is crucial that these advancements are accessible to a broad audience, not limited by specific hardware or software.

In alignment with these technological strides, we present the *Mesoscale Explorer*, an innovative web application designed for exploring large molecular models, spanning from viruses to cell organelles. Primarily targeted at researchers, educators, and students in molecular and structural biology, it offers an interactive experience with guided tours and animations to navigate the complexities of these models. These guided tours are also accessible to a wider general public, providing an opportunity for anyone interested to gain unprecedented access and insight into the molecular fabric of life. Built upon the robust open source *Mol\** framework^39^, *Mesoscale Explorer* leverages cutting-edge web technologies to render intricate details of large-scale models consisting of billions of atoms.

Applications like *Mesoscale Explorer* are at the forefront of the effort to democratize molecular and cellular knowledge, offering intuitive interfaces and guided explorations that demystify the molecular universe.

Accessible online at https://molstar.org/me/, the *Mesoscale Explorer* invites users to delve into the world of molecular landscape visualization. The dedicated landing page offers links to directly access and explore various models and guided tours within the viewer. Additionally, comprehensive documentation and tutorials are available at https://molstar.org/me-docs/, designed to help users maximize their experience with the *Mesoscale Explorer*, ensuring a smooth and enriching journey through the molecular landscapes that underpin life itself.

## Methods

*Mesoscale Explorer* is built on top of *Mol\**, a cutting-edge library designed for developing web applications to visualize molecular data. This initiative, born from a collaboration between *PDBe* and *RCSB PDB* (the European, American branches of the Worldwide Protein Data Bank (https://www.wwpdb.org) and *CEITEC* (https://www.ceitec.eu/), aims to synergize the capabilities of *LiteMol*^18^ (developed by PDBe^40^) and *NGL*^17^ (developed by RCSB PDB^41^) viewers. *Mol\** stands out as a custom rendering engine optimized for molecular graphics on the web and is currently the default interactive viewer at the *RCSB PDB* (www.rcsb.org). In this work, we extend *Mol\** to support real-time rendering of molecular scenes encompassing billions of atoms. To enhance performance and tackle common bottlenecks associated with rendering large-scale scenes, we developed several optimizations inspired by the state of the art implementation introduced by *cellVIEW*^37^.

### Supported Models

Our intention is to support models coming from any available modeling and experimental methods that provide data in a compatible format. We have currently limited our supported format to mmCIF and explored a new generic container, known as a manifest. This manifest, implemented as a ZIP archive, embeds a JSON file that organizes metadata and references associated binary files, allowing for flexible and structured data storage..

#### mmCIF and BinaryCIF

The macromolecular Crystallographic Information File (mmCIF) format and its binary version^42^ developed by the wwPDB^41^ has become the standard in the structural biology domain. This format augments the CIF (Crystallographic Information File) format by intricately detailing the complexity inherent to biological macromolecules, including proteins, nucleic acids, and their complex assemblies. Structured as a text file, mmCIF organizes information into data blocks filled with tagged items, making it intelligible to both humans and computational tools. For expansive models, it employs Biological Assembly Descriptions (e.g., _pdbx_struct_assembly, _pdbx_struct_assembly_gen, _pdbx_struct_assembly_prop, _pdbx_struct_oper_list) to spatially arrange molecule instances within three-dimensional spaces, linking these assemblies to specific protein chains of the entities described. We introduce an optional customization feature, allowing the incorporation of comprehensive markdown descriptions for every entity within the mmCIF schema. Furthermore, our system accommodates the specialized format of *YASARA* Pet-World models^30^, which is an adapted version of mmCIF. These files encapsulate multiple, distinct objects, delineated by progressively increasing atom_site.pdbx_PDB_model_num. The first pdbx_struct_assembly stores their locations, while a *YASARA*-specific extension appends the model number (pdbx_struct_assembly_gen.PDB_model_num) and annotates object names (pdbx_model.name) along with instance numbers (pdbx_model.instances). Our platform also supports markdown descriptions within the *YASARA* mmCIF files, provided they are manually included in the designated fields.

#### Manifest Files

In response to the need for enhanced efficiency in processing generic mesoscale models, we have developed a container format with a manifest file. This format is predicated on a JSON-based structure that meticulously outlines the scene, incorporating a group filter, a comprehensive list of entities (in binary cif format), their group attributes, and spatial information (position and rotation) stored in streamlined binary files. This approach is designed to significantly reduce storage demands while maintaining precision and fidelity in model representation and interaction.

#### *Mol\** session

These files are a dedicated *Mol\** format that captures states and assets of a given session. This format is used to store guided tours and pre-made scenes.

### Reducing Rendering Costs

Real time rendering of billions of atoms is achieved by: 1) taking advantage of patterns in mesoscale models and 2) applying graphics techniques for rendering large scenes from, for example, video games. Our approach includes a) use of instancing to draw multiple copies of identical structures, b) representation of structures with simple sphere geometries, c) reduced detail based on the distance to the camera, d) drawing of only what is potentially visible, that is, not covered and within the camera view. Additionally, we can decrease the rendering resolution and approximate sphere geometries.

For mesoscale models, we assume that all structures of molecular entities of the same type are generally identical. For example, all copies of hexokinase in a cytoplasm model are assumed to have the same structure, meaning they share the same general atomic coordinates. This assumption enables the use of a very fast instancing approach for visualization. Only a single set of atomic coordinates is required for each molecular entity, which can be passed to the GPU once and then used to render all instances of that entity in the mesoscale model by applying their individual positions and rotations. Thus, the atom coordinates can be passed only once to the GPU as a molecular entity and together with a set of positions and rotations many instances of it can be drawn. We use spheres to represent structures because they are a common, familiar type of representation; they can be rendered very efficiently with high quality using ray-casted impostors^43,44^ and they allow easy simplification by combining multiple adjacent spheres into a single sphere to reduce visual clutter and rendering cost. As structures move further from the camera, reducing their visual detail allows us to scale from virus capsid models to representations of small bacteria or cell organelles at a high frame rate. Since mesoscale models are generally densely packed, we can cull (i.e., not draw) structures that are not within the camera (e.g., when zoomed-in to part of the model) or completely covered by nearer structures using frustum or occlusion culling, respectively.

Variable graphical preset modes are available to suit the user’s needs and hardware capabilities. Level of detail (LOD), resolution, and sphere approximation (e.g. flat disc option) are the parameters that vary per mode. As illustrated in **Figure 1**, we currently provide four modes, sorted by decreasing resolution and details, that enable increasingly higher framerate. The **Ultra** mode provides the most detailed view of the model by keeping more atomistic details with very high LOD; the default **Quality** mode provides the best compromise between atomic details with high LOD and framerate; the **Balanced** modes uses sphere approximation and medium LOD; the **Performance** mode focuses on high frame rate at the price of a lower resolution with low LOD.

**Figure 1.**
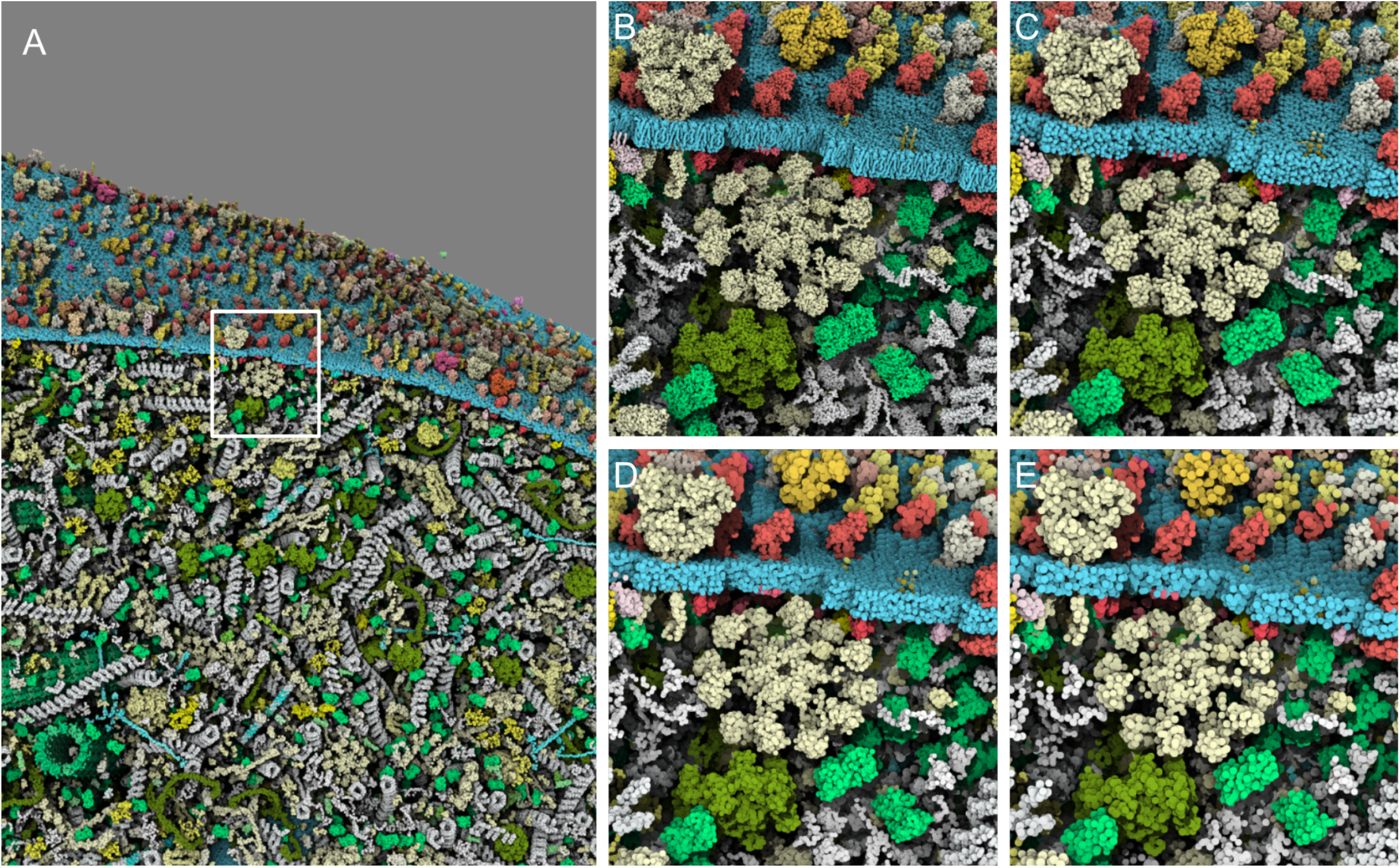
**User-definable options** for quality of rendering with closeup zoom: A. Default quality large view of the model with a white square selection for close-up view: B highly detailed but costly Ultra option, C default detailed rendering Quality mode, D medium rendering quality rendering with approximate sphere Balanced mode and E the lower resolution and highly interactive Performance option. [Explore this interactively](https://molstar.org/me/viewer/?url=https://mesoscope.scripps.edu/explorer/examples/Figure1.molx&type=molx&hide-controls=1).

### Enhancing Perception

Visual perception in molecular rendering can be improved by refining the lighting and shadowing techniques employed. Shadowing can clarify complex 3D spatial relationships and topologies, but traditional shadow rendering often necessitates the computationally intensive task of redrawing the entire scene from the light source’s viewpoint. To circumvent this challenge and streamline the rendering process, we have adopted a screen space shadow technique^45,46^, also known as contact shadowing. This method calculates shadows directly within the screen space by tracing a path from each pixel toward the light source. At each step along this path, we evaluate the depth of our ray against the depth perceived by the camera. If the ray’s depth exceeds that of the camera’s view, it implies that the pixel lies in shadow.

We complement this shadowing technique with Screen Space Ambient Occlusion (SSAO), a revered technique in computer graphics designed to approximate the ambient occlusion effect in real-time. SSAO enhances the depth and realism of 3D scenes by mimicking the way light radiates in real-world environments, particularly how it struggles to penetrate tight spaces. Given the unique demands of molecular landscapes, we tailor our SSAO implementation to suit. Based on the state-of-the-art method from Filion and McNaughton^47^, our customized approach varies the occlusion radius across multiple steps, allowing for a nuanced distinction between close-contact atoms and the broader spatial relationships among molecular chains or protein complexes. As illustrated in **Figure 2**, this dual strategy of tailored shadowing and SSAO aids in the visual differentiation of structural details, facilitating a deeper understanding of complex molecular formations, such as the nuanced interior of a virus capsid.

**Figure 2.**
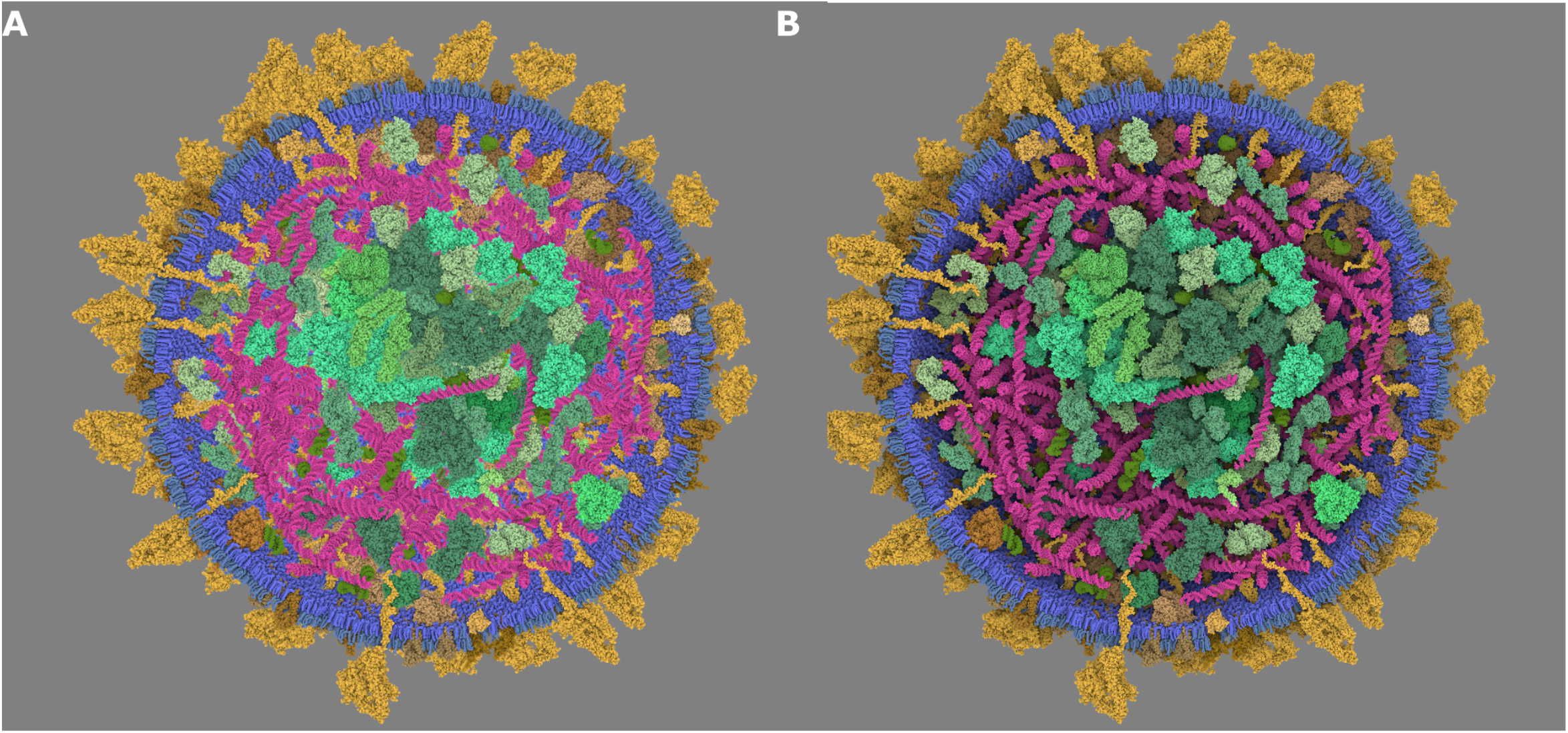
Illustration of the SSAO post-process. **A** - On the left, standard SSAO that emphasizes local occlusion. **B** - On the right, our multiscale approach that captures occlusion at a farther distance. See it interactively at [this link](https://molstar.org/me/viewer/?url=https://mesoscope.scripps.edu/explorer/examples/Figure2.molx&type=molx&hide-controls=1).

By leveraging these advanced lighting techniques, our goal is to not only improve the aesthetic quality of our molecular renderings but also enhance the user’s ability to perceive and interpret intricate molecular structures with greater clarity and insight.

### Clipping

To optimize the interactive visualization and exploration of large molecular landscapes, we have deployed sophisticated clipping strategies (see **Figure 3**). These strategies enable precise control over the parts of the molecular assembly that are visible within the viewport, effectively managing the rendering workload. Our toolkit supports an array of clipping objects based on well-known distance functions and their usage^48^, including planes, spheres, cubes, cylinders, and infinite cones, offering a versatile range of geometric constraints for visual exploration.

**Figure 3.**
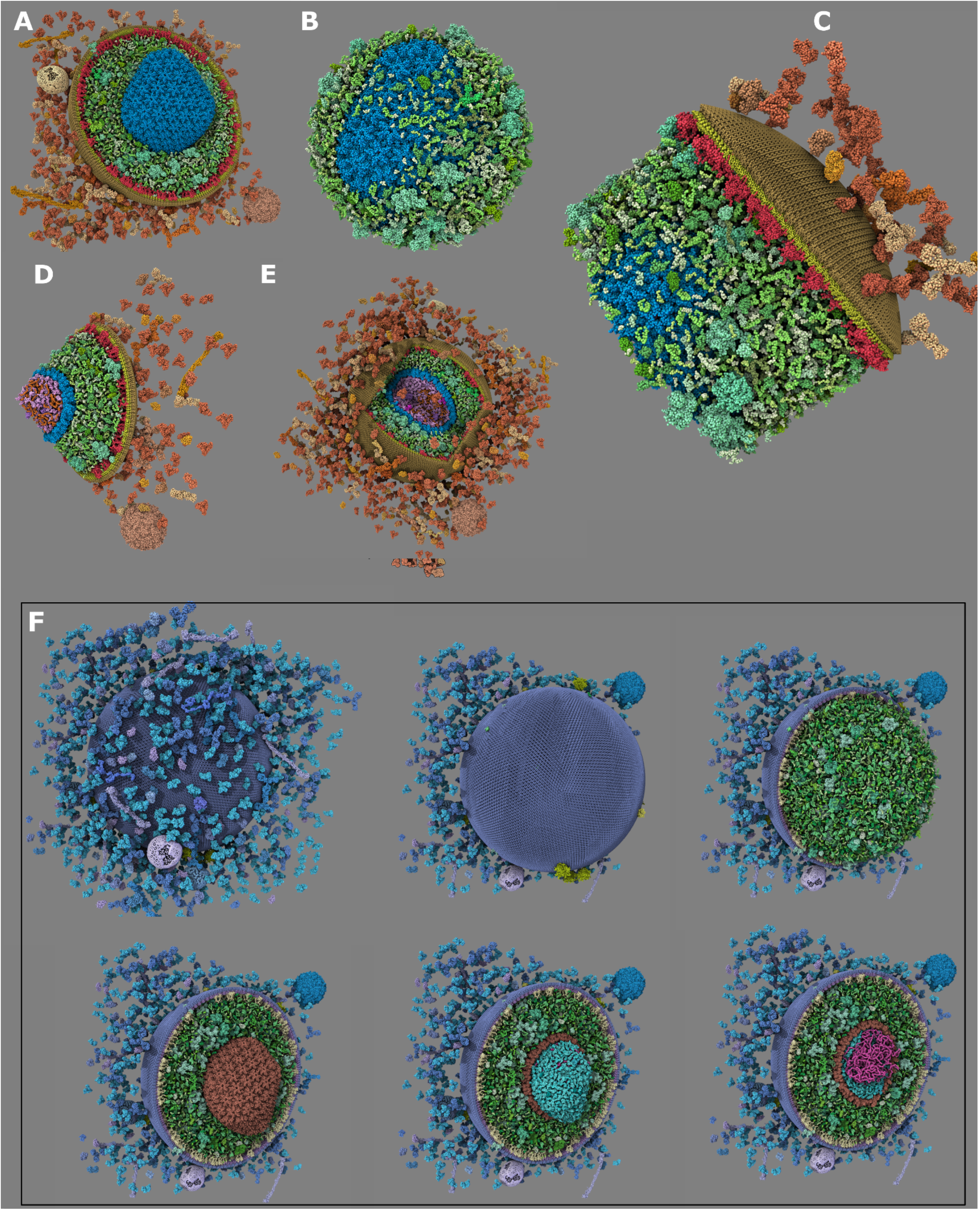
Clipping objects. **A**. plane, **B** sphere (inverted), C cylinder (inverted), D **infinite cone (inverted), and E** cube. **F** Illustrate how we can use progressive clipping to reveal the different features of the HIV structural model, peeling layer by layer to reveal the genome inside. [Explore this interactively](https://molstar.org/me/viewer/?url=https://mesoscope.scripps.edu/explorer/examples/Figure3.molx&type=molx&hide-controls=1).

The power of these clipping mechanisms lies in their selective applicability to distinct subsets of molecular entities. This targeted approach grants researchers the capability to tailor their visual analysis, enabling the crafting of complex visual narratives within the molecular landscape. For example, in **Figure 3F**, a virus model is dissected by clipping planes to expose its internal structure, while simultaneously ensuring that specific elements, such as its genetic material or capsid, remain highlighted and fully visible. This level of detail and flexibility not only enriches the visual representation but also deepens the analytical value of the rendered scenes, allowing for a more nuanced exploration of molecular structures.

Enhancing the visibility of specific protein groups within a model can also be achieved through strategic use of transparency. However, the challenge lies in maintaining visual clarity when the rendered elements are exclusively spheres. Addressing this, we have designed a custom transparency shading to dynamically adjust the transparency, or alpha value, of each sphere based on its normal’s orientation relative to the camera. This means that a sphere becomes more transparent when its normal points away from the user’s perspective. This nuanced approach ensures that transparency not only serves its functional role in altering visibility but also contributes to the overall aesthetic quality of the rendering as illustrated in **Figure 4**.

**Figure 4.**
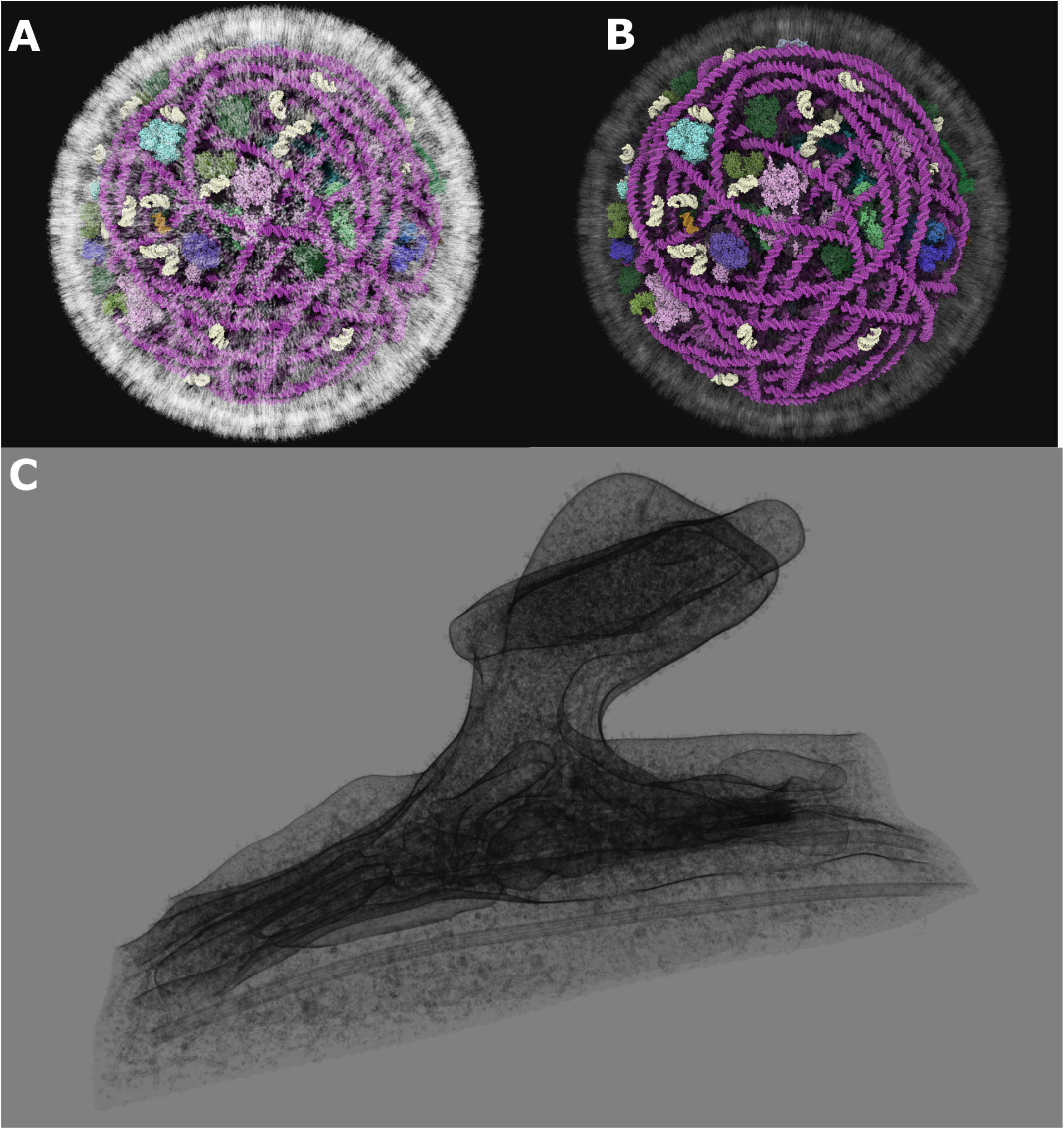
Transparent material. (**A&B**) Applied to the Exosome lipid membrane with two different opacity values (left 0.6 and right 0.1). See it interactively at https://molstar.org/me/viewer/?url=https://mesoscope.scripps.edu/explorer/examples/Figure4AB.molx&type=molx&hide-controls=1. **C**. Transparency (0.015) and black uniform color applied to the whole post-synapse model, revealing its internal structure. [Explore this interactively](https://molstar.org/me/viewer/?url=https://mesoscope.scripps.edu/explorer/examples/Figure4C.molx&type=molx&hide-controls=1).

### Exploration

*Mol\** provides a large range of functionalities that enable users to delve into molecular models via both guided and user-built tours and animations. At the heart of this feature are two foundational *Mol\** modules: snapshots and labeling. Snapshots capture the current state of the viewer and all the parameters currently in use (camera position, clipping, colors, visibility, etc.) stored together with a name, key and description. Labels are 3D text that can be added to any selection. Multiple snapshots can be created at any given moment and re-played akin to a slideshow. Notably, *Mol\** facilitates automatic view interpolation between snapshots, adding a seamless, narrative flow to the presentation. This capability empowers users to weave a story through scenes that highlight and elaborate on significant areas within the models.

The storytelling is enriched through key bindings and Markdown^49^ annotations available in the snapshot description and labels, providing a layered, descriptive narrative over the visualization. Markdown is a lightweight markup language with plain-text formatting syntax. In Markdown, the link text is enclosed in square brackets []. This is the visible text that the reader will click on. The target URL is enclosed in parentheses (). We currently support additional target links for snapshots with a given key, highlighting proteins with a given name, highlighting groups, thereby offering an interactive and immersive exploration tool. **Figure 5** illustrates the rendering of one snapshot description (**Figure 5A**) in a widget viewport and the effect of the mouse hovering on the word ‘proteins’ (**Figure 5B**) that will highlight all the interior proteins of the model.

**Figure 5.**
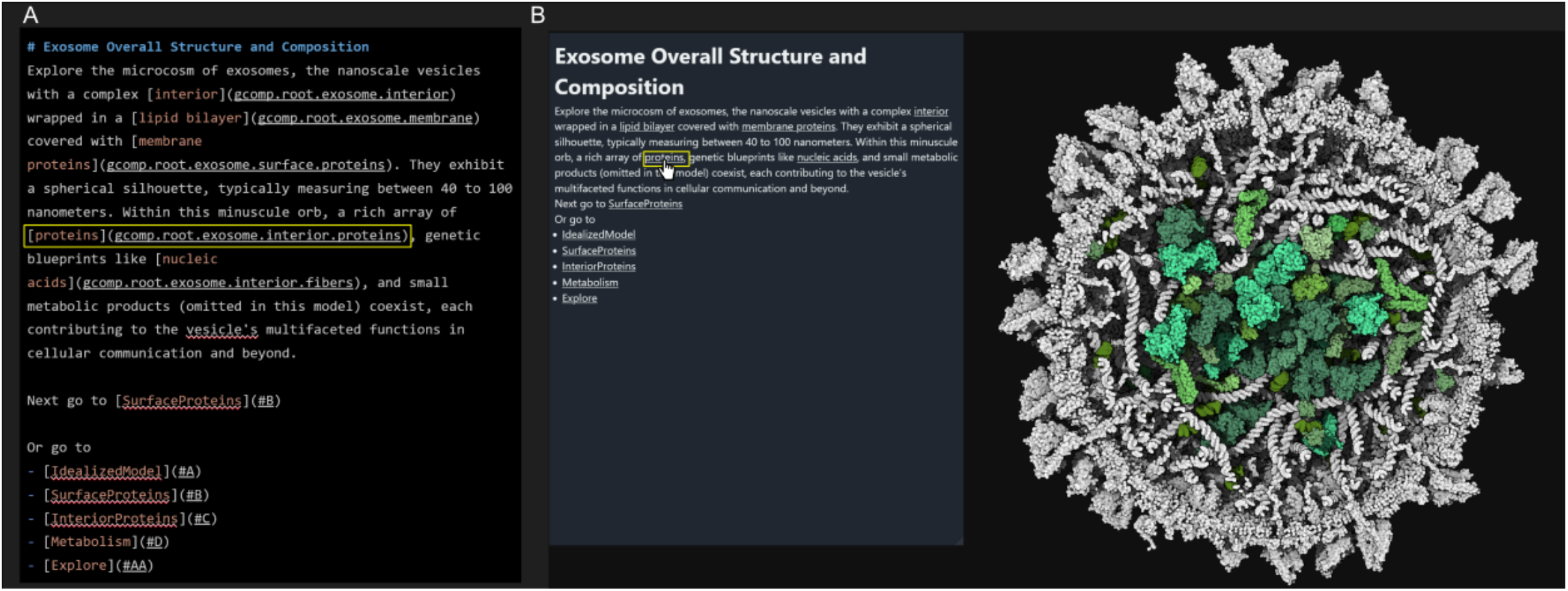
Markdown Description. **A** - Raw description utilizing Markdown syntax to include link to other snapshots as well as mouse hover highlighting feature. **B** - Html rendering of the description and illustration of the mouse hover effect, the mouse is hovering the word ‘protein’ which highlights in the 3d view the interior protein, here in green color, while the rest of the model remains uncolored. [Explore this interactively](https://molstar.org/me/viewer/?example=cellpack-exosome-tour&hide-controls=1).

To further enrich the visualization, 3D labels serve as a powerful tool for annotating selected elements within a scene. These labels, when incorporated into the scene, are not just static markers; they are fully interactive and can be linked directly to snapshot keys. This feature significantly enriches the user experience by making the navigation through molecular tours more intuitive and captivating. Customization includes the label’s text, color, a markdown tooltip for additional information that appears on mouse hover, a reference to a snapshot key that makes the label interactable in the viewer for a quick navigation to the specified snapshot, font style, border appearance, and tethering mechanism. This array of options empowers users to create a personalized and informative visual exploration environment. A complete tutorial on how to make a guided tour is available in the documentation page https://molstar.org/me-docs/tutorial/.

### Guided Tour

We have created tours for many currently published and accessible mesoscale models, including but not limited to the HIV mature virus^30,50^, Mycoplasma genitalium^51^, and Exosomes^52^. A consistent methodology was applied across all tours to ensure a cohesive educational experience. Each journey begins with an introductory snapshot that presents the model’s complete structure. Subsequent snapshots delve deeper, crafting a narrative that highlights features like membrane, critical proteins or biological process, guiding users from the general architecture down to the molecular intricacies.

As users navigate the model, interactive elements come to the forefront. A hover-over feature activates when the mouse cursor moves over individual proteins, dynamically highlighting them and displaying their names in a tooltip window positioned at the bottom of the screen. If a detailed description is available within the model file, this information is also shown, providing valuable context and insights.

The textual content enriching these tours is the product of extensive research, drawn from a wealth of resources including literature reviews, the Protein Data Bank^41^ (*PDB*), *UniProt*^53^, *KEGG*^54^, and *PDB-101*^55^. This information ensures that each tour is grounded in accurate and up-to-date scientific knowledge, offering a rich, immersive learning experience that spans the breadth and depth of mesoscale molecular biology.

## Results & Discussion

To illustrate performance and capabilities of the *Mesoscale Explorer* we provide a landing page with examples of standalone models as well as guided tours. In the sections below, we use standalone models to benchmark the viewer performance, and several guided tours to illustrate usage of the app. **Figure 6** includes some of the currently available models and their corresponding tours, as more fully described in the documentation (https://molstar.org/me-docs/).

**Figure 6.**
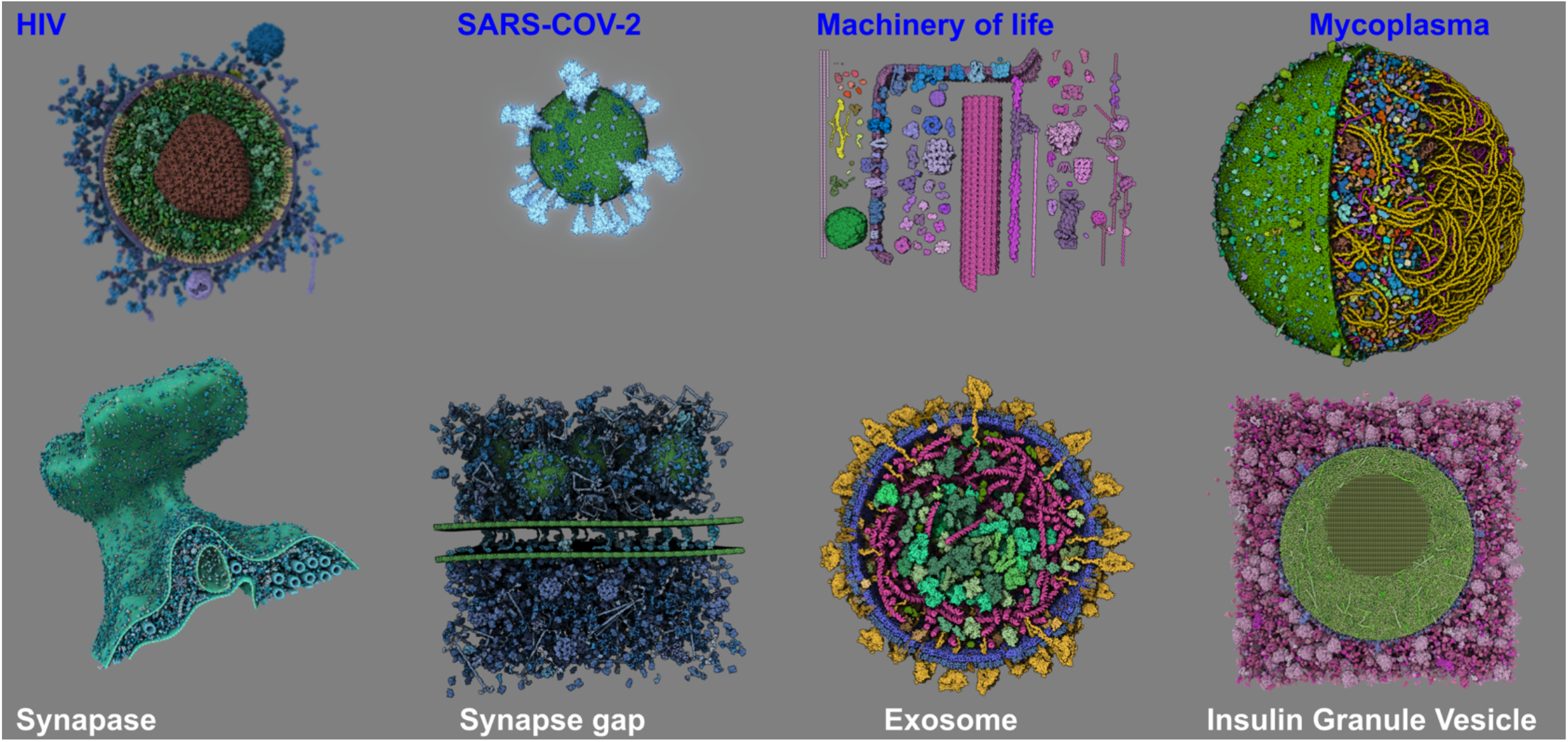
**Examples of some of the** growing number of available models in *Mesoscale Explorer* https://molstar.org/me/. Models with names in blue have guided tours available.

### Model Performance

To evaluate the rendering performance of *Mesoscale Explorer*, we conducted a series of tests on models with different complexity. **Figure 7** presents a histogram of GPU rendering performance, measured in milliseconds (ms), for each model. Lower values on the y-axis indicate better rendering performance. The default ‘Quality’ setting emerged as the best compromise between performance and visual quality, achieving interactive performance for all models, even those with up to 3 billion atoms. Rendering performance was measured from a default view, which maintains a consistent distance from the model, as rendering performance is dependent on the camera’s proximity. The Level of Detail (LOD) feature ensures that performance improves as the camera moves further from the object, while close-up views are more demanding on the GPU.

**Figure 7.**
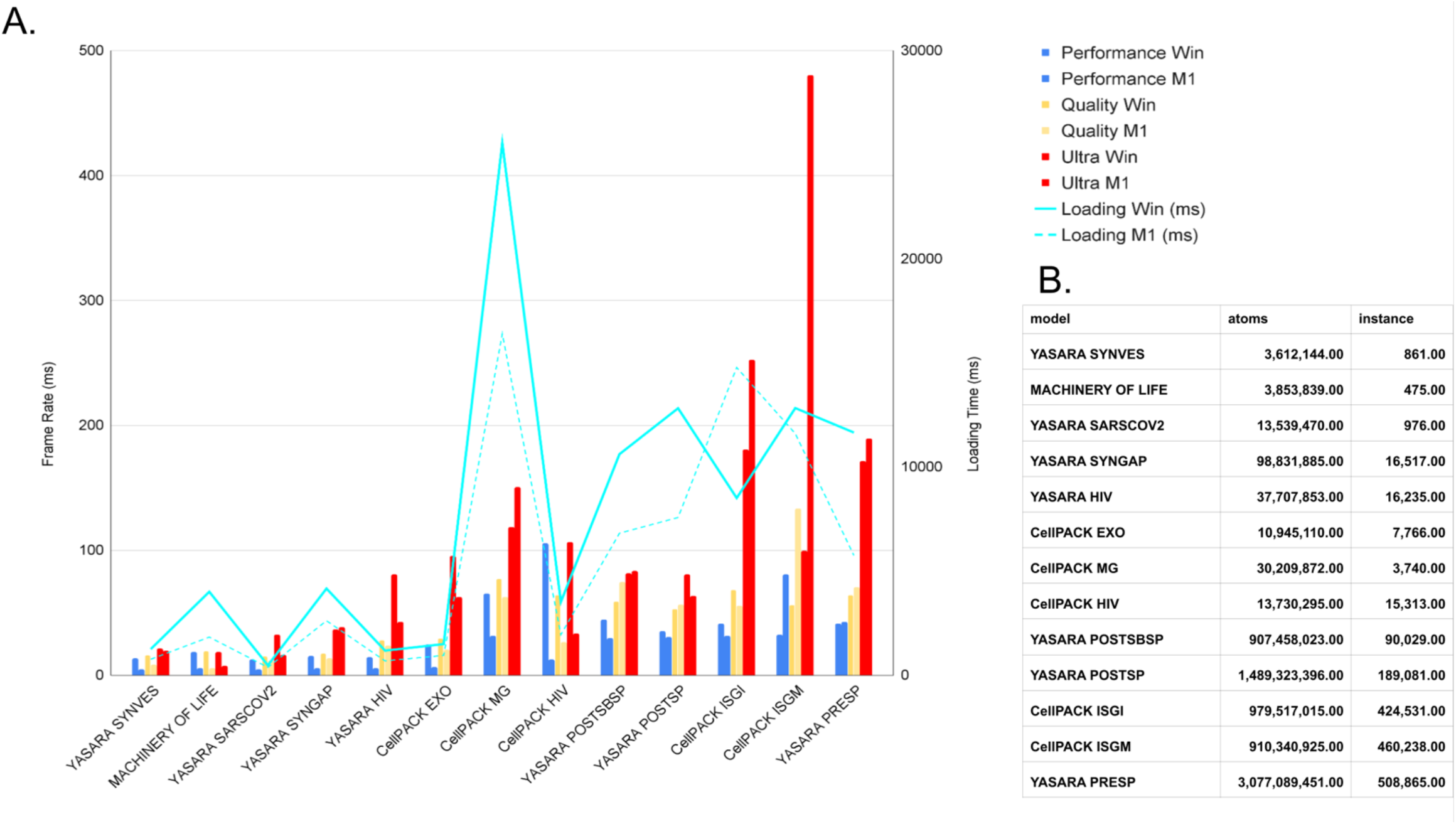
Average Performance. **A**. Histogram of GPU rendering performance (left vertical axis in ms) combined with loading time (right vertical axis in ms) for various example models. The measurements were taken from the default view after loading each model and applying the rock animation. The x-axis shows the model names, and the y-axis displays the performance (lower values indicate better performance). Each color represents a quality setting (Quality, Performance, Ultra) and a machine (Windows, *Win*, with AMD Threadripper 1950X and Quadro RTX 8000 or Mac OS with apple silicon *M1*) at a resolution of 1598×1776. **B**. Table of number of atoms and instances for each mode.

### Tours Overview

Starting from a storyboard draft, the time required to produce a tour is approximately 2 hours. The time needed to acquire the necessary information is not easily quantifiable, as it represents the culmination of years of research. However, the process of gathering and summarizing this information is streamlined through web-based interfaces like *UniProt*^53^, as well as pre-existing materials from *PDB-101*^55,56^ and *Wikipedia*. While we have attempted to utilize current Large Language Models (LLMs) to enhance our research, the results have been suboptimal. The information provided currently by LLMs often lacks sufficient detail, contains excess information, or includes inaccuracies and requires manual curation. Below we will describe some of the tours we developed to highlight some useful features offered by Mesoscale Explorer.

### Virus: SARS-COV-2 and HIV

SARS-CoV-2 is arguably the most recognizable virus for most of the general public since the pandemic.

We used one of the currently available models developed by the *YASARA* Team^30^ and plan to support alternative models made with the Mesocraft^31^ modeling method. Being one of the smallest models, this tour, illustrated in **Figure 8A and 8B**, is easily run on any device compared to the larger billions-atom models such as the pre-synapse model.

**Figure 8.**
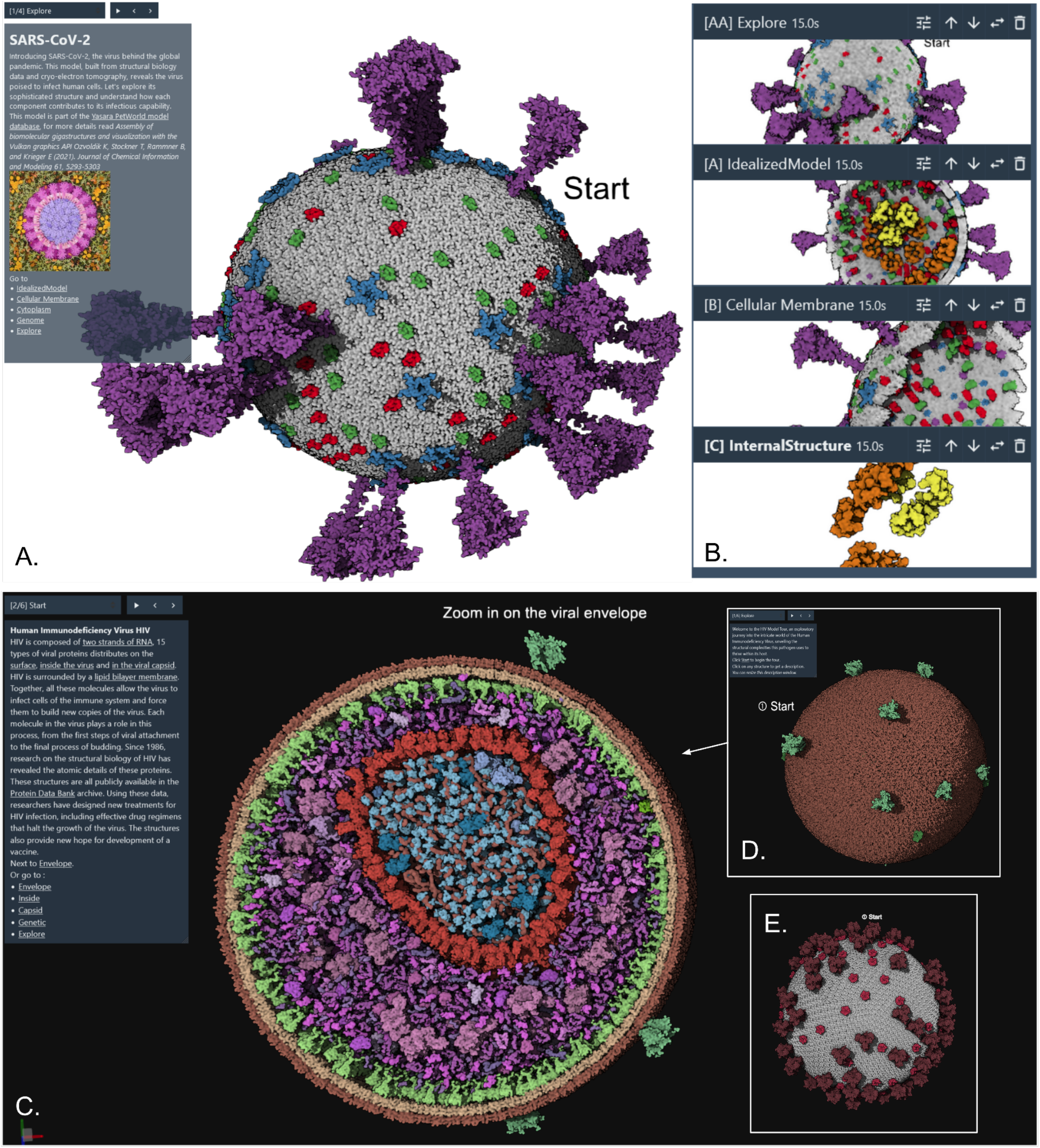
Tours of Viruses. **A**. Initial default view showing an overview of the entire SARS-CoV-2 virion. **B**. Four snapshots from the [tour](https://molstar.org/me/viewer/?example=petworld-sarscov2-tour&hide-controls=1), exploring structural aspects of the envelope and interior. **C**. *cellPACK* HIV tour showing interior view and **D** starting snapshot. **E**. *YASARA* PetWorld HIV tour starting snapshot.

In addition, two models of mature HIV virions are currently available to the community, one from the *YASARA* PetWorld database^30^ and one from the *cellPACK* team^50^ (see **Figure 8 C,D,E**). *Mesoscale Explorer* allows exploration of the overall similarity of the ultrastructure and components of both models, and highlights differences between them. For example, the *YASARA* model includes two genomic RNA of the virion using known secondary structure, while the *cellPACK* model is a random walk model of the two single strand gRNAs. Users can also explore differences in the distribution of Matrix protein under the lipid membrane, as well as the lipids membrane itself. *YASARA* uses a rhombus tiling approach while *cellPACK* uses a modified lipidWrapper^57^ approach designed to run on GPU.

### Bacteria: Mycoplasma

This tour is the longest, because of its complexity and the availability of a great amount of metadata.

Every entry has *UniProt*^53^ and *KEGG*^54^ information. For instance, we can retrieve a pathway and design a markdown description that uses ASCII art to provide a graphical map of a reaction pathway. As shown in **Figure 9**, we use this approach to showcase the glycolysis pathway.

**Figure 9.**
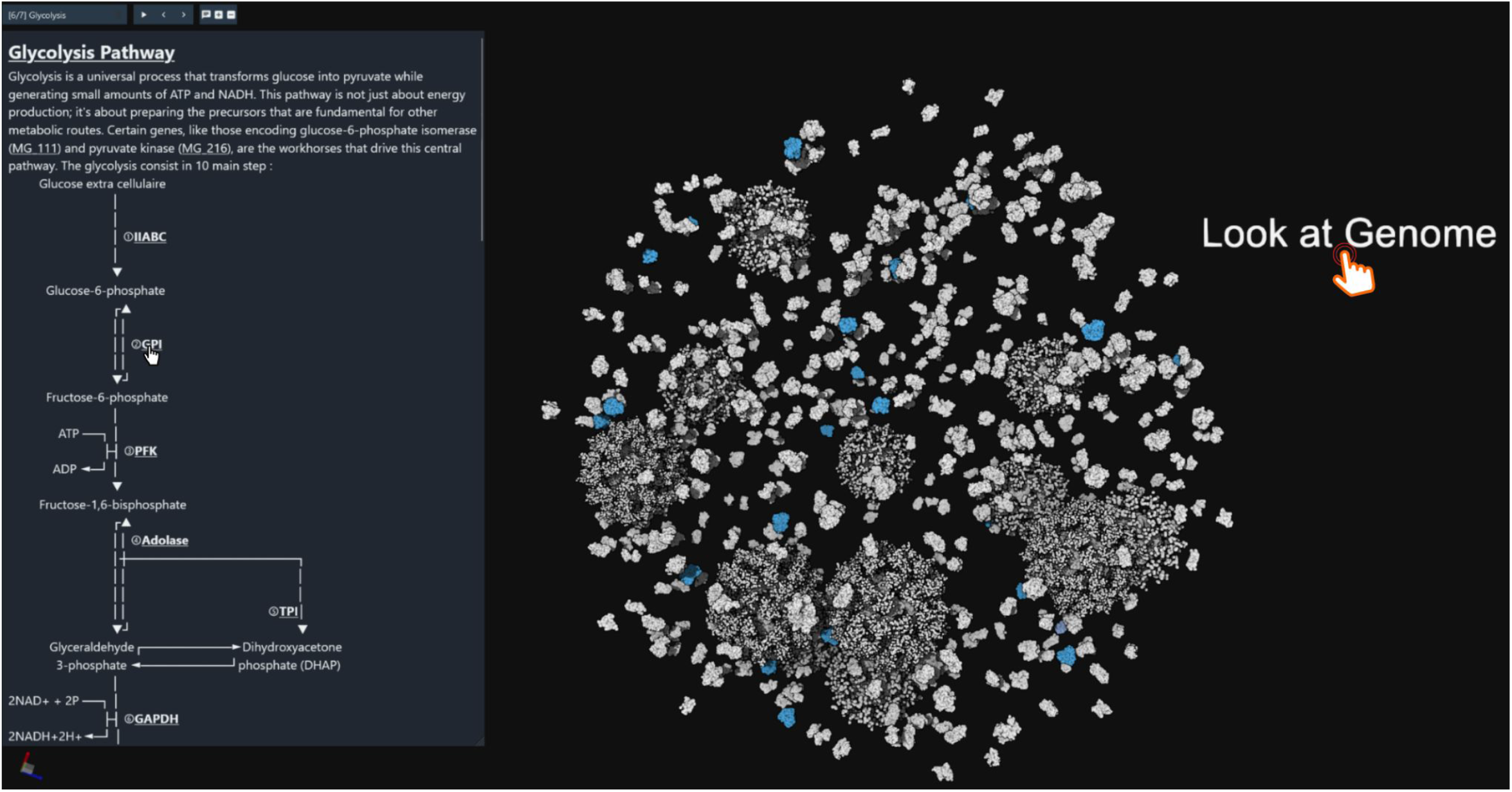
Mycoplasma tour example of guided snapshot of the glycolysis pathway. The description shows the pathway, with mouse over effects that highlight in the model the underlying enzyme. An interactable label can be seen, ‘Look at Genome’, which will bring the user to the next snapshot if clicked.

### Machinery of life

In 2002, the *RCSB PDB* published “Molecular Machinery: A Tour of the Protein Data Bank.” This illustration, available as a poster and handout, has been distributed to thousands of PDB users, teachers, and students over the years. To celebrate the release of over 100,000 structures in the PDB archive in 2014, the RCSB PDB updated this iconic image in a new edition titled “Tour of the Protein Data Bank.” This new edition is also available as a 2d interactive poster (**Figure 10A**). We ported it to *Mesoscale Explorer* and created a 3D interactive tour of the poster (**Figure 10B**). The vast range of molecular shapes and sizes in the PDB is illustrated by 96 molecular machines. The structures are depicted relative to the cellular membrane and organized into categories related to function. Using the poster, we manually placed and oriented all the given PDB IDs and grouped them using the same classification. Each snapshot corresponds to a given category and provides a labeled view of all the proteins forming the group (**Figure 10C and 10D**).

**Figure 10.**
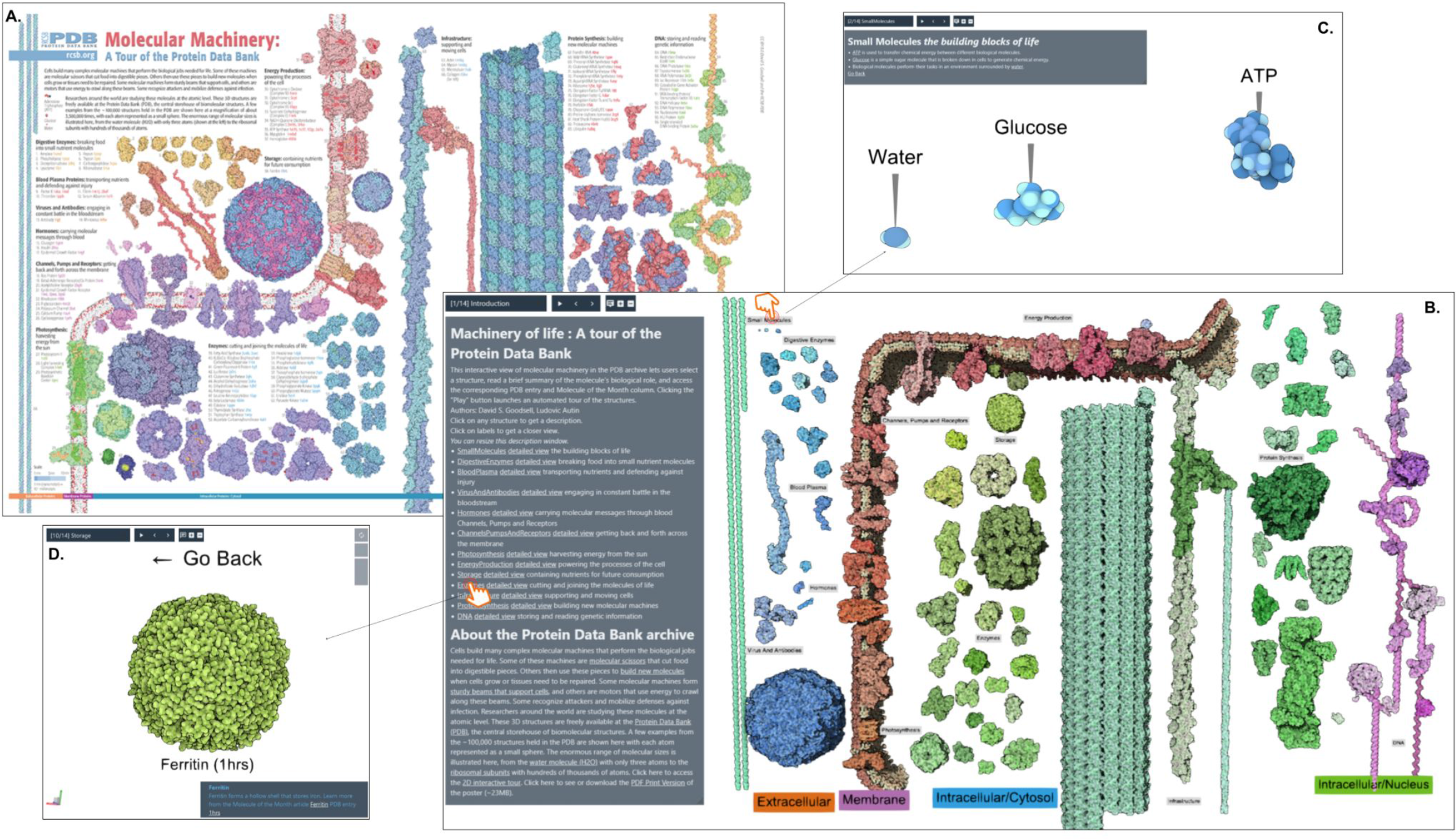
**Machinery of Life tour**, exploring diverse biomolecules from the Protein Data Bank archive. **A**. Poster available at RCSB Protein Data Bank website. **B**. Interactive version created in *Mesoscale Explorer*. **C**. Detailed views of individual molecules are provided by clicking on the overall view. **D**. Detailed views of the Storage category provided by clicking on the description hyperlink. [Explore this interactively](https://molstar.org/me/viewer/?example=machineryoflife-tour&hide-controls=1).

### FUTURE DIRECTION

*Mol\** is an open-source project that is actively being developed, ensuring that the *Mesoscale Explorer* will benefit from all advancements in the core library. Moreover, now that the foundational elements are in place, we are poised to support additional file formats; to explore automated tours inspired by Molecumentary^32,58^, to implement better story editing inspired by ScrollyVis^59^; to develop enhancements in advanced lighting effects such as global illumination; as well as supporting dynamic data similar to the *Simularium Viewer*^60^.

## CONCLUSION

In our endeavor, we’ve struck a critical balance between rendering fidelity and efficiency by implementing strategies like level of detail (LOD) and culling. Contrary to intuition, reducing rendering fidelity can enhance perceptual clarity, minimizing visual noise and directing focus towards pivotal details. This optimization, coupled with the guided tour functionality, elevates the user experience by weaving a narrative through the intricate molecular terrains, facilitating interactive engagement with these complex structures.

The widespread accessibility of molecular visualization tools, accelerated by the COVID-19 pandemic, has opened new avenues for researchers, educators, and the broader public to engage with molecular science. Platforms like Mesoscale Explorer, built upon the well-tested framework of Mol*, are at the forefront of this democratization effort. The foundational data management and visualization features of Mol* are robust and stable, allowing ready prototyping of features such as LOD rendering and efficient rendering of group of spheres instances that are specific to mesoscale models and guided tours. Features developed for Mesoscale Explorer can be used in Mol*, and vice versa, ensuring a cohesive and flexible development environment.

The absence of a standardized format or repository for mesoscale models and their guided tours highlights the need for a unified system. Initiatives like MolViewSpec^61^ propose a standard format that could be extended for such tours, while PDB-DEV^62^ offers a potential repository for storing these models. The creation and maintenance of these standards will likely involve community-driven efforts, focusing on licensing, contribution strategies, and iterative refinement processes.

Fueled by the exponential growth in computational power and rendering technologies, the domain of molecular visualization is advancing rapidly. This progress is not only deepening our understanding of the molecular underpinnings of life but also setting the stage for groundbreaking discoveries and applications across medicine, biotechnology, and related fields. As we continue to peel back the layers of biological complexity, we move closer to unveiling the intricate web of processes that orchestrate life itself.

## Acknowledgements

The authors gratefully acknowledge input on design of the tours from Pr. Arthur Olson. This work was supported in part by grants GM120604 and 5U54AI170855 from the National Institutes of Health (DSG and LA), and a grant from the Czech Science Foundation [22-30571M] (DS).

## Data Availability

WebApp release and its documentation: https://molstar.org/me/. Open-source repo on Github: https://github.com/molstar/molstar

## Author contributions

A.R. developed the app, D.S contributed to the app development; D.G. contributed to the guided tour design and L.A. did project management, contributed to the app development, and authored the tours. All authors contributed to writing the manuscript.

## REFERENCES

1. Levinthal C (1966) Molecular model-building by computer. Sci. Am. 214:42–52.

2. Marshall GR, Beitch J, Ellis RA, Fritsch JM (1972) Macromolecular modeling system: the insulin dimer. Diabetes 21:506–508.

3. McGill G (2008) Molecular Movies… Coming to a Lecture near You. Cell 133:1127–1132.

4. Riggi M, Torrez RM, Iwasa JH (2024) 3D animation as a tool for integrative modeling of dynamic molecular mechanisms. Struct. Lond. Engl. 1993 32:122–130.

5. O’Donnell TJ, Olson AJ (1981) GRAMPS - A graphics language interpreter for real-time, interactive, three-dimensional picture editing and animation. ACM SIGGRAPH Comput. Graph. 15:133–142.

6. Richardson DC, Richardson JS (1992) The kinemage: a tool for scientific communication. Protein Sci. Publ. Protein Soc. 1:3–9.

7. Sayle R (1995) RASMOL: biomolecular graphics for all. Trends Biochem. Sci. 20:374–376.

8. Humphrey W, Dalke A, Schulten K (1996) VMD: Visual molecular dynamics. J. Mol. Graph. 14:33–38.

9. Carson M (1997) Ribbons. Methods Enzymol. 277:493–505.

10. Sanner MF (1999) Python: a programming language for software integration and development. J. Mol. Graph. Model. 17:57–61.

11. Delano WL The PyMOL Molecular Graphics System (2002). In: ; 2002. Available from: https://api.semanticscholar.org/CorpusID:203708320

12. Moll A, Hildebrandt A, Lenhof H-P, Kohlbacher O (2005) BALLView: an object-oriented molecular visualization and modeling framework. J. Comput. Aided Mol. Des. 19:791–800.

13. Pettersen EF, Goddard TD, Huang CC, Couch GS, Greenblatt DM, Meng EC, Ferrin TE (2004) UCSF Chimera—A visualization system for exploratory research and analysis. J. Comput. Chem. 25:1605–1612.

14. Pettersen EF, Goddard TD, Huang CC, Meng EC, Couch GS, Croll TI, Morris JH, Ferrin TE (2021) UCSF ChimeraX : Structure visualization for researchers, educators, and developers. Protein Sci. 30:70–82.

15. Ozvoldik K, Stockner T, Krieger E (2023) YASARA Model–Interactive Molecular Modeling from Two Dimensions to Virtual Realities. J. Chem. Inf. Model. 63:6177–6182.

16. Anon Jmol: an open-source Java viewer for chemical structures in 3D. Available from: http://www.jmol.org/

17. Rose AS, Bradley AR, Valasatava Y, Duarte JM, Prlic A, Rose PW (2018) NGL viewer: web-based molecular graphics for large complexes. Bioinforma. Oxf. Engl. 34:3755–3758.

18. Sehnal D, Deshpande M, Vařeková RS, Mir S, Berka K, Midlik A, Pravda L, Velankar S, Koča J (2017) LiteMol suite: interactive web-based visualization of large-scale macromolecular structure data. Nat. Methods 14:1121–1122.

19. Wang J, Youkharibache P, Zhang D, Lanczycki CJ, Geer RC, Madej T, Phan L, Ward M, Lu S, Marchler GH, et al. (2020) iCn3D, a web-based 3D viewer for sharing 1D/2D/3D representations of biomolecular structures. Bioinforma. Oxf. Engl. 36:131–135.

20. Hodis E, Schreiber G, Rother K, Sussman JL (2007) eMovie: a storyboard-based tool for making molecular movies. Trends Biochem. Sci. 32:199–204.

21. Goddard T (2017) Creating animations with UCSF ChimeraX: spin, morph, density fit, and virtual reality movies. Available from: https://www.rbvi.ucsf.edu/chimera/data/wcpcw-mar2017/moviemaking.html

22. Johnson GT, Autin L, Goodsell DS, Sanner MF, Olson AJ (2011) ePMV Embeds Molecular Modeling into Professional Animation Software Environments. Structure 19:293–303.

23. Porollo A, Meller J (2010) POLYVIEW-MM: web-based platform for animation and analysis of molecular simulations. Nucleic Acids Res. 38:W662–W666.

24. Maiti R, Van Domselaar GH, Wishart DS (2005) MovieMaker: a web server for rapid rendering of protein motions and interactions. Nucleic Acids Res. 33:W358–362.

25. Autin L, Tufféry P (2007) PMG: online generation of high-quality molecular pictures and storyboarded animations. Nucleic Acids Res. 35:W483–488.

26. Raush E, Totrov M, Marsden BD, Abagyan R (2009) A New Method for Publishing Three-Dimensional Content Gay N, editor. PLoS ONE 4:e7394.

27. Kühlbrandt W (2014) The Resolution Revolution. Science 343:1443–1444.

28. Johnson GT, Autin L, Al-Alusi M, Goodsell DS, Sanner MF, Olson AJ (2015) cellPACK: a virtual mesoscope to model and visualize structural systems biology. Nat. Methods 12:85–91.

29. Klein T, Autin L, Kozlikova B, Goodsell DS, Olson A, Groller ME, Viola I (2018) Instant Construction and Visualization of Crowded Biological Environments. IEEE Trans. Vis. Comput. Graph. 24:862–872.

30. Ozvoldik K, Stockner T, Rammner B, Krieger E (2021) Assembly of Biomolecular Gigastructures and Visualization with the Vulkan Graphics API. J. Chem. Inf. Model. 61:5293–5303.

31. Nguyen N, Strnad O, Klein T, Luo D, Alharbi R, Wonka P, Maritan M, Mindek P, Autin L, Goodsell DS, et al. (2021) Modeling in the Time of COVID-19: Statistical and Rule-based Mesoscale Models. IEEE Trans. Vis. Comput. Graph. 27:722–732.

32. Kouril D, Strnad O, Mindek P, Halladjian S, Isenberg T, Groller ME, Viola I (2023) Molecumentary: Adaptable Narrated Documentaries Using Molecular Visualization. IEEE Trans. Vis. Comput. Graph. 29:1733–1747.

33. Kadir SR, Lilja A, Gunn N, Strong C, Hughes RT, Bailey BJ, Rae J, Parton RG, McGhee J (2021) Nanoscape, a data-driven 3D real-time interactive virtual cell environment. eLife 10:e64047.

34. Alharbi R, Strnad O, Luidolt LR, Waldner M, Kouril D, Bohak C, Klein T, Groller E, Viola I (2023) Nanotilus: Generator of Immersive Guided-Tours in Crowded 3D Environments. IEEE Trans. Vis. Comput. Graph. 29:1860–1875.

35. Lindow N, Baum D, Hege H -C. (2012) Interactive Rendering of Materials and Biological Structures on Atomic and Nanoscopic Scale. Comput. Graph. Forum 31:1325–1334.

36. Falk M, Krone M, Ertl T (2013) Atomistic Visualization of Mesoscopic Whole-Cell Simulations Using Ray-Casted Instancing. Comput. Graph. Forum 32:195–206.

37. Le Muzic M, Autin L, Parulek J, Viola I (2015) cellVIEW: a Tool for Illustrative and Multi-Scale Rendering of Large Biomolecular Datasets. Eurographics Workshop Vis. Comput. Biomed. 2015:61–70.

38. Alharbi R, Strnad O, Hadwiger M, Viola I (2024) Nanouniverse: Virtual Instancing of Structural Detail and Adaptive Shell Mapping. Available from: http://arxiv.org/abs/2404.05116

39. Sehnal D, Bittrich S, Deshpande M, Svobodová R, Berka K, Bazgier V, Velankar S, Burley SK, Koča J, Rose AS (2021) Mol* Viewer: modern web app for 3D visualization and analysis of large biomolecular structures. Nucleic Acids Res. 49:W431–W437.

40. Varadi M, Anyango S, Appasamy SD, Armstrong D, Bage M, Berrisford J, Choudhary P, Bertoni D, Deshpande M, Leines GD, et al. (2022) PDBe and PDBe-KB: Providing high-quality, up-to-date and integrated resources of macromolecular structures to support basic and applied research and education. Protein Sci. Publ. Protein Soc. 31:e4439.

41. Berman HM (2000) The Protein Data Bank. Nucleic Acids Res. 28:235–242.

42. Sehnal D, Bittrich S, Velankar S, Koča J, Svobodová R, Burley SK, Rose AS (2020) BinaryCIF and CIFTools-Lightweight, efficient and extensible macromolecular data management. PLoS Comput. Biol. 16:e1008247.

43. Gumhold S Splatting Illuminated Ellipsoids with Depth Correction. In: International Symposium on Vision, Modeling, and Visualization. ; 2003. Available from: https://api.semanticscholar.org/CorpusID:5889109

44. Grottel S, Reina G, Ertl T Optimized data transfer for time-dependent, GPU-based glyphs. In: 2009 IEEE Pacific Visualization Symposium. Beijing, China: IEEE; 2009. pp. 65–72. Available from: http://ieeexplore.ieee.org/document/4906839/

45. Karabelas P (2020) Screen space shadows. Available from: https://panoskarabelas.com/posts/screen_space_shadows/

46. Aldridge G (2023) Screen Space Shadows. Available from: https://s2023.siggraph.org/presentation/?id=exs104&sess=sess437

47. Filion D, McNaughton R Effects & techniques. In: ACM SIGGRAPH 2008 Games. Los Angeles California: ACM; 2008. pp. 133–164. Available from: 10.1145/1404435.1404441

48. Quilez I (2008) Modeling with distance functions. Model. Distance Funct. [Internet]. Available from: https://iquilezles.org/articles/distfunctions/

49. Gruber J (2004) Daring Fireball. Markdown [Internet]. Available from: https://daringfireball.net/projects/markdown/

50. Johnson GT, Goodsell DS, Autin L, Forli S, Sanner MF, Olson AJ (2014) 3D molecular models of whole HIV-1 virions generated with cellPACK. Faraday Discuss 169:23–44.

51. Maritan M, Autin L, Karr J, Covert MW, Olson AJ, Goodsell DS (2022) Building Structural Models of a Whole Mycoplasma Cell. J. Mol. Biol. 434:167351.

52. Jiménez J, Autin L, Ibáñez de Cáceres I, Goodsell DS (2019) Integrative Modeling and Visualization of Exosomes. J. Biocommun. 43:e10.

53. UniProt Consortium (2023) UniProt: the Universal Protein Knowledgebase in 2023. Nucleic Acids Res. 51:D523–D531.

54. Kanehisa M, Furumichi M, Sato Y, Kawashima M, Ishiguro-Watanabe M (2023) KEGG for taxonomy-based analysis of pathways and genomes. Nucleic Acids Res. 51:D587–D592.

55. Zardecki C, Dutta S, Goodsell DS, Lowe R, Voigt M, Burley SK (2022) PDB-101: Educational resources supporting molecular explorations through biology and medicine. Protein Sci. Publ. Protein Soc. 31:129–140.

56. Goodsell DS, Dutta S, Zardecki C, Voigt M, Berman HM, Burley SK (2015) The RCSB PDB “Molecule of the Month”: Inspiring a Molecular View of Biology. PLOS Biol. 13:e1002140.

57. Durrant JD, Amaro RE (2014) LipidWrapper: an algorithm for generating large-scale membrane models of arbitrary geometry. PLoS Comput. Biol. 10:e1003720.

58. Kouril D, Isenberg T, Kozlikova B, Meyer M, Groller ME, Viola I (2021) HyperLabels: Browsing of Dense and Hierarchical Molecular 3D Models. IEEE Trans. Vis. Comput. Graph. 27:3493–3504.

59. Mörth E, Bruckner S, Smit NN (2023) ScrollyVis: Interactive Visual Authoring of Guided Dynamic Narratives for Scientific Scrollytelling. IEEE Trans. Vis. Comput. Graph. 29:5165–5177.

60. Lyons B, Isaac E, Choi NH, Do TP, Domingus J, Iwasa J, Leonard A, Riel-Mehan M, Rodgers E, Schaefbauer L, et al. (2022) The Simularium Viewer: an interactive online tool for sharing spatiotemporal biological models. Nat. Methods 19:513–515.

61. Bittrich S, Midlik A, Varadi M, Velankar S, Burley SK, Young JY, Sehnal D, Vallat B (2024) Describing and Sharing Molecular Visualizations Using the MolViewSpec Toolkit. Curr. Protoc. 4:e1099.

62. Vallat B, Webb B, Fayazi M, Voinea S, Tangmunarunkit H, Ganesan SJ, Lawson CL, Westbrook JD, Kesselman C, Sali A, et al. (2021) New system for archiving integrative structures. Acta Crystallogr. Sect. Struct. Biol. 77:1486–1496.

